# Uncoupling the CRMP2-Ca_V_2.2 interaction reduces pain-like behavior in a preclinical osteoarthritis model

**DOI:** 10.1101/2024.06.05.596514

**Authors:** Heather N. Allen, Sara Hestehave, Paz Duran, Tyler S. Nelson, Rajesh Khanna

**Author notes:** To whom correspondence should be addressed: Dr. Rajesh Khanna, Department of Pharmacology & Therapeutics, and Pain and Addiction Therapeutics (PATH) Collaboratory, University of Florida College of Medicine, Gainesville, Florida 32610. These authors contributed equally (Heather N. Allen, Sara Hestehave). Department of Experimental Medicine, University of Copenhagen, 2200 Copenhagen N, Denmark (Sara Hestehave).

## Abstract

Osteoarthritis (OA) represents a significant pain challenge globally, as current treatments are limited and come with substantial and adverse side effects. Voltage-gated calcium channels have proved to be pharmacologically effective targets, with multiple FDA-approved CaV2.2 modulators available for the treatment of pain. Although effective, drugs targeting CaV2.2 are complicated by the same obstacles facing other pain therapeutics-invasive routes of administration, narrow therapeutic windows, side effects, and addiction potential. We have identified a key regulator of CaV2.2 channels, collapsing response mediator protein 2 (CRMP2), that allows us to indirectly regulate CaV2.2 expression and function. We developed a peptidomimetic modulator of CRMP2, CBD3063, that effectively reverses neuropathic and inflammatory pain without negative side effects by reducing membrane expression of CaV2.2. Using a rodent model of OA, we demonstrate the intraperitoneal administration of CBD3063 alleviates both evoked and non-evoked behavioral hallmarks of OA pain. Further, we reveal that CBD3063 reduces OA-induced increased neural activity in the parabrachial nucleus, a key supraspinal site modulating the pain experience. Together, these studies suggest CBD3063 is an effective analgesic for OA pain.

## Introduction

Chronic pain represents a significant health concern globally, imposing substantial economic burdens on healthcare systems and affecting the quality of life for millions of individuals^15^. It is estimated that chronic pain affects over 1.5 billion people worldwide, leading to an annual economic toll of hundreds of billions of dollars due to medical costs, lost productivity, and disability claims^15,43,48^. Among the various forms of chronic pain, osteoarthritis (OA) stands out as a prevalent and debilitating condition^35^.

OA, characterized by the progressive degeneration of joint cartilage and underlying bone, affects approximately 10% of men and 18% of women aged 60 years and older^51^. This degenerative joint disease is one of the leading causes of disability among older adults, significantly impairing mobility and quality of life^47^. The increasing prevalence of OA is attributed to an aging population, obesity, and joint injuries, necessitating effective management strategies to alleviate pain and improve function^37^.

Current pharmacological treatments for OA pain include nonsteroidal anti-inflammatory drugs (NSAIDs), acetaminophen, and opioids^16^. While these medications provide symptomatic relief, they are often associated with significant adverse effects. NSAIDs, though effective in reducing inflammation and pain, can cause gastrointestinal ulcers, cardiovascular issues, and renal impairment with long-term use^1,46^. Acetaminophen, commonly used for mild to moderate pain, may lead to hepatotoxicity in high doses^13^. Opioids, though potent analgesics, carry the risk of addiction, tolerance, and a wide array of side effects, including respiratory depression and constipation^4^. Given these limitations, there is a pressing need for new therapeutic agents that can effectively manage OA pain with minimal adverse effects.

Transmembrane CaV2.2 (N-type) voltage-gated calcium channels are genetically and pharmacologically validated pain targets^27^. Three commercially available analgesics target CaV2.2: Ziconotide (Prialt®), gabapentin (Neurontin®), and Pregabalin (Lyrica®).

Unfortunately, the use of these drugs is hindered for numerous reasons, including invasive routes of administration, narrow therapeutic windows, sedation, and abuse liabilities^20,30,42^. Nevertheless, preclinical^29,50^ and clinical^19,45^ data demonstrate the efficacy of targeting CaV2.2 for alleviating arthritis-associated pain. Therefore, improvements in targeting CaV2.2 may represent a promising approach for treating OA pain.

We previously found that the microtubule-binding collapsin response mediator protein 2 (CRMP2) is a key regulator of CaV2.2 trafficking and function^9,11^. Overexpression of CRMP2 in sensory neurons enhanced CaV2.2 currents, surface expression, and neurotransmitter release^11^. Additionally, we identified a 15 amino-acid peptide (designated CBD3, for calcium channel binding domain 3) that effectively interferes with the CaV2.2–CRMP2 interaction, decreasing calcium influx, neurotransmitter release, and inflammatory and neuropathic pain^7^. More recently, our team identified a CBD3-based peptidomimetic, designated CBD3063, that uncoupled CaV2.2 from CRMP2, reduced membrane CaV2.2 expression and Ca^2+^ currents, decreased neurotransmission, and reversed neuropathic and inflammatory pain without changes in sensory, sedative, depressive, nor cognitive behaviors^24^. Here, we test the hypothesis that CBD3063 will be efficacious in alleviating pain-like behavior in the preclinical rodent monoiodoacetate (MIA) model of OA pain. Our data suggest that this compound may offer a promising alternative to current pain management options, potentially addressing the unmet need for safer and more effective OA therapies.

## Methods

### Animals and housing

#### Rats

Male and female pathogen-free Sprague Dawley rats (Charles River Laboratories, Wilmington MA) were used for the behavioral experiments. All animals were pair-housed in transparent IVC cages (37*25*18 cm), in light (12-h light: 12-h dark cycle; lights on at 07:00 h) and temperature (23 ± 3°C) controlled rooms. Each cage was provided with environmental enrichment consisting of paper-wool shavings (Enviropack) for nesting material, Nylabones, and cardboard tube for hiding, and cages were changed twice a week, but never on the days of testing. Food (Purina lab Diet 5053) and water were available *ad libitum* with water changed on a weekly basis. Animals were 6 weeks old upon arrival and were left to acclimatized to the surroundings for at least 1 week before the start of any experiments.

#### Mice

Male and female C57BL/6 mice (The Jackson Laboratory, Bar Harbor, ME) were kept in light (12-h light: 12-h dark cycle; lights on at 07:00 h) and temperature (23 ± 3°C) controlled rooms, with free access to food and water and weekly cage-change. Mice were housed in groups of 4-5 mice and provided with nestlets for nesting material. Animals were 8-10 weeks of age at experiment initiation.

All animal use was conducted in accordance with the Guide for the Care and Use of Laboratory Animals, published by the National Institutes of Health guidelines, and the study was conducted with approval from the New York University Institutional Animal Care and Use Committee (IACUC). All efforts were made to minimize animal suffering.

#### Drug administration

For *in vivo* use, (3R)−3−acetamido−N−[3−(pyridin−2− ylamino)propyl]piperidine−1−carboxamide (designated hereafter as CBD3063) (10 mg/kg)^24^, gabapentin (30 mg/kg)^24^, and vehicle (10% DMSO, 90% saline) was administered via the intraperitoneal (i.p.) route in a volume of 5 ml/kg for rats and 10 ml/kg for mice.

### MIA induction

Induction of the monoiodoacetate arthritis model to the knee joint (MIA) was performed as previously reported by our group^26^. On the day of injection, the monoiodoacetate solution was freshly prepared in sterile saline for injection of Sodium Iodoacetate Bioultra (Sigma, I9148-5G) in saline. The injection-technique was similar for rats and mice however, dose and volume were different for each species (rats; 2 mg/50 μl, mice; 1 mg/10 μl). Animals were anesthetized in an induction chamber using Isoflurane 2.5% mixed in O2 at a flowrate of 1.5 L/min and maintained via facemask at 2.0% during the injection. Anesthetic depth was confirmed by lack of withdrawal reflex to a pinch to the tail.

The animal was then placed on the back in dorsal recumbency, and the fur was shaved in the area around the knee on the left hindleg. The knee was stabilized and fixed in a slightly bent position and the patellar tendon was visualized as a white line below the skin. Using a 30G insulin-syringe (BD Micro-Fine Plus Demi, 0.3 ml) the injection was made intraarticularly in the joint-space by applying it perpendicularly through the skin and tendon just below the patella.

#### Behavioral testing

All behavioral experiments were performed by female experimenters blinded to treatment.

### Mechanical allodynia, VF

Mechanical sensitivity was assessed using a series of calibrated von Frey monofilaments (North Coast Medical, Inc., Morgan Hill), as previously described^26^. The MIA-injury was induced unilaterally in the left knee, but as traditionally performed, the sensory changes (vF, ADT) are not measured at the primary site of injury (the knee-joint), but on the plantar surface of the ipsilateral paw. The increased sensitivity detected via this method is thereby measured in a referred receptive field area and reflects secondary hyperalgesia, indicative of central sensitization^40^. Clinical studies in knee-OA patients show direct correlation between clinical pain experience and the level of sensitization at referred locations^3^.

Animals were placed in individual Plexiglas (9.5*14*19.0 cm) enclosures on an elevated wire grid. They were given 15-60 minutes to acclimate to the enclosure and the experimenter’s presence and movements below the grid. Next the plantar surface of the hind paw was stimulated with a series of calibrated von Frey filaments (0.4, 0.6, 1.0, 1.4, 2.0, 4.0, 6.0, 8.0, 10.0, 15.0, 26.0 g). To initiate testing a filament with a bending force of 4.0 g was first applied to the hind paw with uniform pressure for 5 seconds. A brisk withdrawal was considered a positive response whereupon the next lower filament in the series was applied. In the absence of a positive response the neighboring higher filament was applied. After the first change in response-pattern, indicating the threshold, 4 additional applications were performed; when no response, the next filament with a higher force was tested, and when positive response, the next lower force filament was tested. The 50% threshold was determined by the following equation: 50% threshold (g) = 10^log(last filament)+k*0.3^. The constant, *k*, was found in the table by Dixon^18^, and determined by the response-pattern. Baselines were recorded before (Pre-MIA) and two weeks after injury (Post-MIA), and at 30 min, 1h, 2h, 3h, 4h, 5h and 6 hours after injection of CBD3063, gabapentin, or vehicle.

### Cold allodynia, ADT

Cold allodynia was assessed as previously reported^26,32^. While the animals were still in the Plexiglas chambers following von Frey measurements, cold allodynia was assessed using application of a drop of acetone (Acetone Drop Test, ADT) to the plantar surface of the paw using a plastic syringe without mechanically touching the skin. Following application, the duration of the response (flinching, licking, stepping, guarding) was then recorded, for a maximum of 30 seconds. The application and assessment was performed two times per animal with 5-10 minutes between each application, and the average of the two measurements was calculated. Baselines were recorded before (Pre-MIA) and two weeks after injury (Post-MIA), and at 2h, 4h and 6hours after injection of CBD3063, gabapentin or vehicle.

### Conditioned Place Aversion, CPA

CPA tests were conducted in rats using a two-chamber device based on the protocols from Zhou et al.^55^, and as previously reported by our group^26^. The protocol includes four consecutive tests of 10 min duration: preconditioning (10 min), conditioning (2*10 min) and testing (10 min). During preconditioning, the animal is allowed free access to two connected chambers (30*30*19 cm), each associated with a scented lip-balm applied to the walls (ChapStick Spearmint or Strawberry Lip-Balm). Immediately following preconditioning, a divider was applied between the chambers, and the rat was conditioned to either stimuli or no-stimuli for 10 min in each chamber. The stimuli consisted of repeated stimulation with a 10 g vF-filament every 30 seconds for the 10 min that the subject was contained in that chamber, while no stimuli (NS) was applied when in the other chamber. The order and side of conditioning was alternated between subjects. Following the conditioning, the divider was removed, and the rat was allowed free access to both chambers for the 10 min test. Animal movements in each chamber was recorded by a camera above and analyzed using the ANY-maze software (Stoelting Co. IL, USA). The distance travelled and the duration of time spent in each chamber was recorded during preconditioning and test. Decreased time spent in a chamber during the test versus preconditioning indicated avoidance for that chamber and was calculated as a CPA-score; time in VF-chamber during preconditioning – time in VF-chamber during test. The CPA experiment was conducted 3 weeks after MIA-injury, and testing was initiated 1h after i.p injection of CBD3063, gabapentin or vehicle.

### Functional impairment - Static Weight Bearing, WB

To assess the functional impact of the injury the static weight bearing distribution was assessed as previously described^26^. Hindlimb weight bearing was measured using a Bioseb Incapacitance Test (Bioseb) which measures the weight distribution across the two hindlimbs of a stationary animal. Animals were habituated and trained to become comfortable with the testing paradigm during short sessions for 2 days prior to induction of the model. Three readings were collected and averaged for each animal, and the weight borne by the ipsilateral limb was expressed as a percentage of the weight borne across both hindlimbs (WB-% = (weight borne on the injured leg/weight borne on both legs) * 100%). Baselines were recorded before (Pre-MIA) and three weeks after injury (Post-MIA), and at 2 hours after injection of CBD3063, gabapentin or vehicle.

#### *In vivo* calcium imaging, fiber photometry, FP

As previously reported by our group^24,26^, adult male and female wildtype mice received 500 nL of AAV9-CaMKIIa-GCaMP6s-WPRE-SV40 (Addgene, Watertown, MA) in the right parabrachial nucleus (PBN) to transfect glutamatergic PBN neurons with the calcium indicator GCaMP6s (coordinates: A/P-5.15 mm, M/L+/-1.45 mm, D/V-3.45 mm). Virus was precisely administered with a Nanoject II Auto-Nanoliter Injector (Drummond) at a rate of 2 nL/sec and a wait time of 5 minutes to prevent backflow. Directly following viral infusion, a fiber optic cannula with black ceramic ferrule (RWD, 1.25 mm ferrule diameter, 200 µm core diameter, and 0.37 numerical aperture) was chronically implanted in the right PBN and fixed to the skull using dental cement (Cat# 10-000-786, Stoelting, Wood Dale, Il). Mice were allowed 21 days to recover before undergoing baseline testing. Mice were acclimated in acrylic boxes on wire mesh with fiber optic patch cord attached for at least one hour prior to testing. Calcium transients were collected continuously (FP3002, Neurophotometrics) during stimulation protocol. The plantar surface of the left hindpaw was exposed to the following stimuli: 1 second application of a 0.07 g and 1.0 g von Frey filament, 10 µL acetone drop, and blunted pinprick. Each stimulus was repeated three times with two minutes between stimuli, and all three responses were averaged to represent the animal’s response. Using custom MatLab scripts, the GCaMP6s signal (470 nm laser) was normalized to the isosbestic control 405 nm laser signal to control for photobleaching and motion artifacts. Change in fluorescence (dF/F) was calculated by subtracting the GCaMP6s signal following stimulation from the average of GCaMP6s signal for 10 seconds prior to stimulation. Area under the curve was calculated for the ten seconds directly following stimulus application. Area under the curves were log transformed for normality. The day after baseline recordings, MIA was induced as described above. Two weeks following MIA, the fiber photometry protocol was repeated in the same animals to collect post MIA responses of glutamatergic neurons in the PBN to mechanical stimuli. The same fiber photometry protocol was conducted again 1-2 hours after intraperitoneal injection of CBD3063 (10 mg/kg), gabapentin (30 mg/kg) and vehicle (10% DMSO). Following the completion of the experiment, animals were transcardially perfused with ice cold 1x PBS and 10% neutral buffered formalin (Cat# SF98-4, Fisher Scientific, Waltham, MA) before brains were extracted for verification of viral infection and fiberoptic placement. 30 µm thick coronal brain sections were obtained on a cryostat and stored at 4°C. To visualize GCaMP6s expression, we performed immunohistochemistry for GFP. Briefly, sections were washed 3 times in PBS for 5 minutes, incubated in normal goat serum (Cat# 5425, Cell Signaling Technology, Danvers, MA) based blocking buffer (PBS with 5% normal goat serum 0.1% Triton X-100) for one hour, and incubated in primary antibody (Rabbit anti-GFP 1:1000 in blocking buffer, Cat# AB3080, Millipore Sigma, St. Louis, MO) overnight at room temperature on an orbital shaker. Sections were then washed 3 times in PBS with 0.1% Triton X-100 for 5 minutes, incubated for 1.5 hours in secondary antibody (goat anti rabbit AlexaFluor 488, Cat# A11008, Invitrogen, Waltham, MA), and washed again in PBS before being mounted on SuperFrost Plus microscope slides (Cat# 22-037-246, Fisher Scientific, Waltham, MA), coverslipped with Vectashield Plus antifade mounting medium with DAPI (H-2000-10, Vector Laboratories), and imaged at 20x on Leica DMi8 inverted widefield microscope. No animals were excluded due to post hoc target verification. Please note that the fiber photometry animals were tested with multiple analgesic compounds spaced in time to conserve the total number of animals used. The pre-MIA, post-MIA, and vehicle data were previously published^26^ and are reproduced in this paper with permission of the publisher.

### Molecular interrogation

#### Synapse enrichment and fractionation

As previously described^26^, naïve and MIA-injured male and female rats were anesthetized with 5% isoflurane and then decapitated. Spinal cords were removed, and the dorsal region of the lumbar spinal cord was dissected as this structure contains the synapses arising from the DRG. Synaptosomes isolation was done as described previously^26^. Integrity of non-postsynaptic density (non-PSD) and PSD fractions was verified by immunoblotting. PSD95 was enriched in the PSD fractions while synaptophysin was enriched in non-PSD fractions.

#### Immunoblot Preparation and Analysis

Indicated samples were loaded on 4–20% Novex gels (Cat# XP04205BOX; Thermo Fisher Scientific). Proteins were transferred for 1 h at 100 V using TGS [25 mM Tris, pH 8.5, 192 mM glycine, 0.1% (mass/vol) SDS], 20% (vol/vol) methanol as transfer buffer to PVDF membranes (0.45 μm; cat. no. IPFL00010; Millipore), preactivated in pure methanol. After transfer, the membranes were blocked at room temperature for 1 h with TBST (50 mM Tris·HCl, pH 7.4, 150 mM NaCl, 0.1% Tween 20) with 5% (mass/vol) nonfat dry milk, and then incubated separately in indicated primary antibodies (1/1,000 dilution), Flotillin 1 (Cat# F1180, RRID:AB_1078893; Sigma-Aldrich, St. Louis, MO), CRMP2 (Cat# C2993, RRID:AB_1078573; Sigma-Aldrich, St. Louis, MO), CaV2.2 (Cat# TA308673, RRID:AB_2650547; Origene, Rockville, MD), NaV1.7 (Cat# ab85015, RRID:AB_2184346; Abcam), PSD-95 (Cat# MA1-045, RRID:AB_325399; Invitrogen), Synaptophysin (Cat# MAB5258, RRID:AB_2313839; Sigma-Aldrich) in TBST, 5% (mass/vol) BSA, overnight at 4 °C. Following incubation in HRP-conjugated secondary antibodies from Jackson Immuno Research (West Grove, PA) (1/10,000 dilution), Mouse Anti-Rabbit (Cat# 211-032-171, RRID:AB_2339149) and Goat Anti-Mouse (Cat# 115-035-174, RRID:AB_2338512), blots were revealed by enhanced luminescence (WBKLS0500; Millipore Sigma St. Louis, MO). Please note, that the immunoblots were run with multiple antibodies (both Nav1.7 and Cav2.2) in the same protein to conserve the number of animals used. Thus, the immunoblot controls CRMP2 and Flotillin 1 were published before^26^ and are reproduced in this paper with permission of the publisher.

#### Data analysis and statistics

Animals were randomly allocated to treatment. Statistical analysis was performed using GraphPad Prism version 9 (GraphPad Software, Inc., La Jolla, CA, USA). Full analysis results are displayed in the supplementary table S1. *P* < 0.05 was considered statistically significant. Error bars in the graphs represents mean ± SEM. For behavioral time-course experiments, the data was also transformed into Area Under the Curve (AUC) for the first 6mhours after injection for each individual animal, in order to simplify treatment comparison.

All details of statistical tests and results are presented for each figure in the **Supplementary Table S1**.

## Results

### MIA induces an increase in CaV2.2 spinal presynaptic localization

In experimental models of chronic pain, elevated levels of Cav2.2 protein have been reported ^14,36,53^. To assess if the monoiodoaceteate (MIA)-injury induces an increase in Cav2.2 spinal presynaptic localization, we used a synaptic fractionation approach. We collected the lumbar dorsal horn from naïve and MIA-injured rats and isolated the pre-synaptic fraction. We found that MIA-injury significantly increases Cav2.2 presynaptic localization in male and female rats (**Fig. 1A-C**). Consistent with our previous data from neuropathic models^33^, no significant difference in total CRMP2 levels was detected between naïve and MIA-injured rats (**Fig. 1A-C**), as previously published^26^.

**Fig 1.**
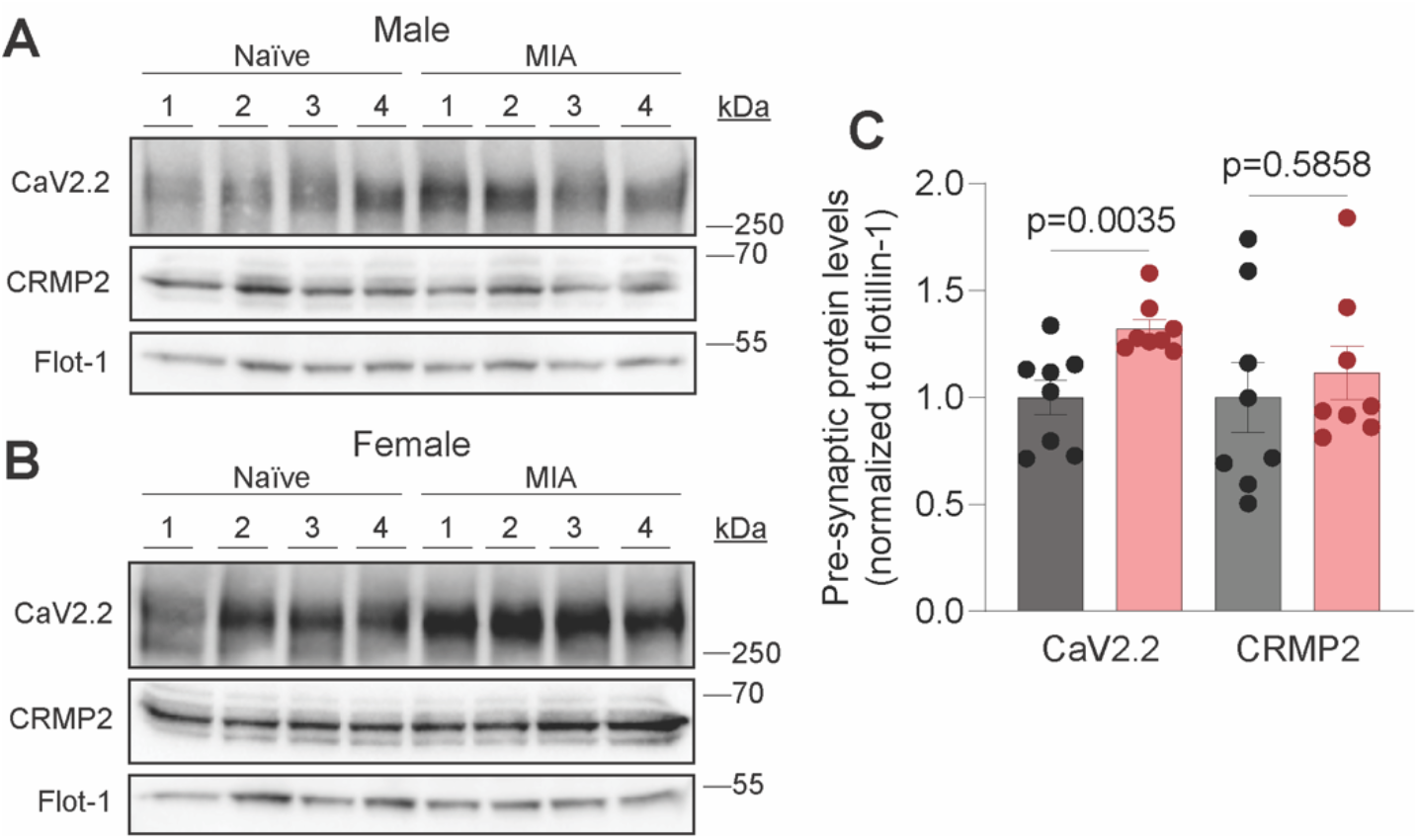
MIA enhances the presynaptic fraction of CaV2.2. Representative immunoblots of presynaptic fractions of lumbar dorsal horn samples from male (A) and female (B) Naïve and MIA-injured rats probed with antibodies against CaV2.2, CRMP2, and membrane-associated protein Flotillin-1 (loading control) (n=8, 4 male/4 female). (C) Bar graph with scatter plots showing quantification of CaV2.2 and CRMP2 in the presynaptic fraction (n=8 rats per condition, 4 male/4 female). Error bars indicate mean ± SEM; data was analyzed with unpaired two-tailed t tests, p values as indicated.

### CBD3063 reduces MIA-induced mechanical and cold allodynia

The MIA model of arthritis was induced by injecting monosodium iodoacetate into the left knee joint of rats. Two weeks later, we compared the anti-hyperalgesic efficacy of CBD3063 (10 mg/kg, i.p.) and gabapentin (30 mg/kg, i.p.) over a 6-hour period post-injection (**Fig. 2A**). Gabapentin was used as a reference compound for two reasons: 1) it has demonstrated anti-hyperalgesic effects in the MIA model^17^, and 2) it alleviates pain by disrupting the Cav2.2–α2d-1 interaction, preventing channel trafficking to the plasma membrane, providing a mechanistically relevant comparator to CBD3063^24^. As previously described^26,39^, MIA reliably induced robust mechanical allodynia in both male and female rats at 2 weeks post-induction, as evidenced by the reduction in mechanical threshold from pre-MIA to post-MIA (**Fig. 2B-C**). CBD30363 significantly reduced MIA-induced mechanical allodynia in both male (**Fig. 2B**) and female (**Fig. 2C**) rats from 1 to 4 hours after injection, whereas gabapentin’s effect lasted from 1 to 3 hours. There were no significant sex-related differences in treatment outcomes across the study-duration (**Fig. 2D**).

**Fig 2.**
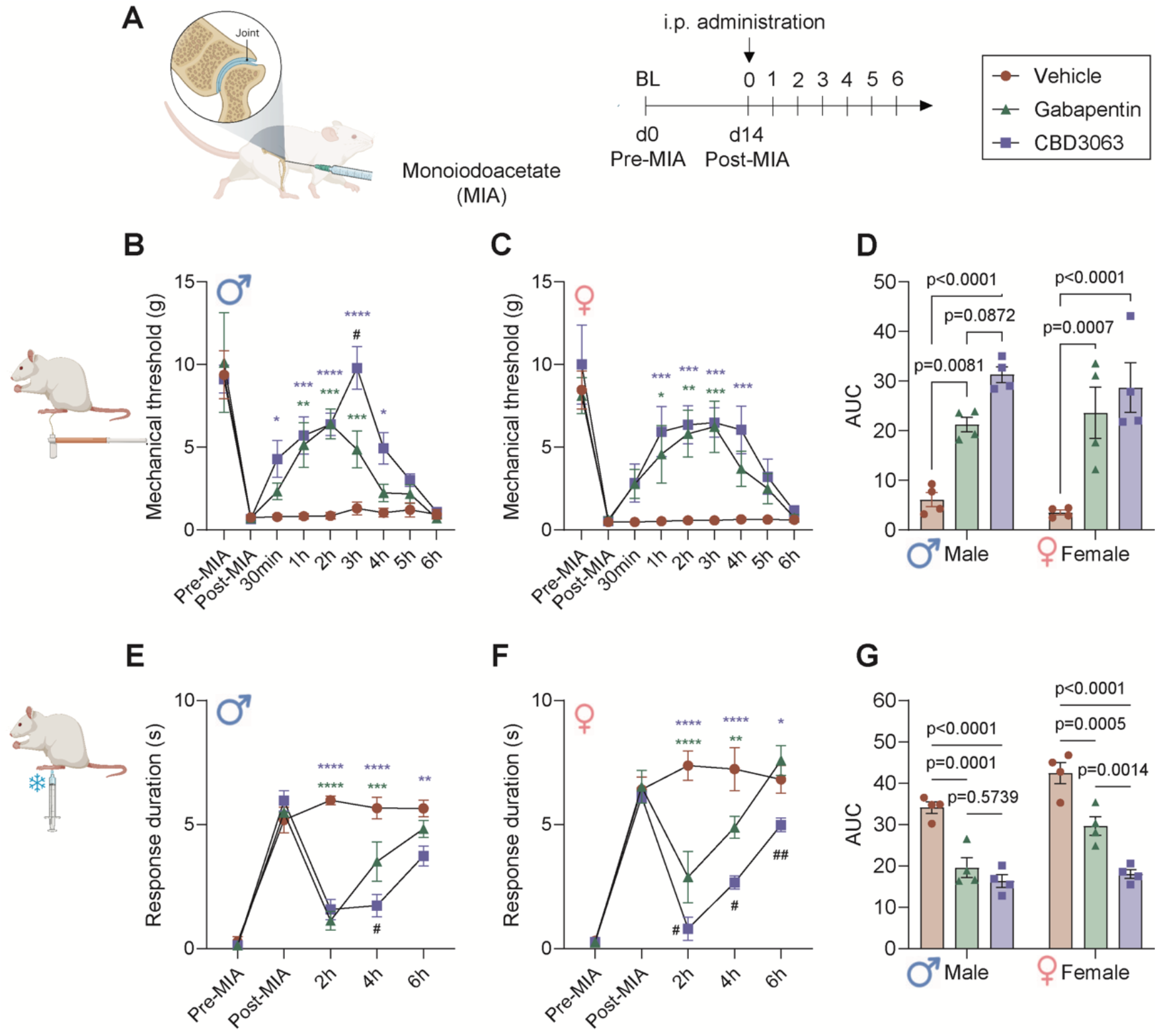
CBD3063 decreases mechanical and cold allodynia in rats with MIA-induced pain-like behaviors. (A) Study-design schematic and treatment conditions. (B-C) Baseline paw-withdrawal threshold was measured before (pre-MIA) and after (post-MIA) in both male (D) and female (C) rats. The analgesic effect of single i.p. injections of CBD3063 (10 mg/kg), gabapentin (30 mg/kg) or vehicle were measured hourly for 6h after administration. n=4 rats pr gender/group. (D) Quantification of Area Under the Curve of the paw withdrawal threshold reported in panel B-C from post-MIA to 6h post administration. Both CBD3063 and Gabapentin significantly improved the MIA-induced mechanical allodynia in both genders. (E-F) Baseline response duration to an application of an acetone drop was measured before (pre-MIA) and after (post-MIA) in both male (E) and female (F) rats. The antiallodynic effect of single i.p. injections of CBD3063 (10 mg/kg), Gabapentin (30 mg/kg) or vehicle were measured for 6h after administration. n=4 rats pr gender/group. (G) Quantification of Area Under the Curve of the response duration reported in panel E-F from post-MIA to 6h post administration. Both CBD3063 and Gabapentin significantly improved the MIA-induced cold allodynia, but CBD3063 was significantly more significant than gabapentin. Results were compared using 2-way ANOVA and post-comparison between groups using Tukey’s multiple comparison test, as suggested by comparison with vehicle-groups; *P<0.05, **P<0.01, ***P<0.001, ****P<0.0001; differences between gabapentin and CBD3063 suggested by, #P<0.05, ##P<0.01. Results are displayed as Mean ± S.E.M.

Similarly, the MIA model induced robust cold allodynia, evidenced by an increased duration of aversive responses to the application of an acetone drop (**Fig. 2E and F**). CBD3063 and gabapentin also reduced the MIA-induced cold allodynia in both male (**Fig. 2E**) and female (**Fig. 2F**) rats. Notably, CBD3063’s analgesic effects were significantly superior to gabapentin’s at 4 hours post-injection in males and throughout the experiment in females. Statistical analysis of the area under the curve (AUC) revealed a significant effect of sex on this outcome measure (**Fig. 2G**), but no significant interaction between sex and treatment. This suggests that the treatment effects were not markedly different between the two sexes, but rather that MIA-injured female rats generally displayed prolonged aversive behavior following cold stimuli compared to males. Post hoc comparison confirmed that CBD3063 was significantly more effective at relieving cold allodynia than gabapentin in females (**Fig 2G**).

### CBD3063 alleviates MIA-induced hyperexcitability of the parabrachial nucleus

Next, we tested if CBD3063 effectively dampened supraspinal pain transmission. The majority of ascending nociceptive information is relayed via the spino-parabrachio-amygdaloid pathway in rodents^34^. Therefore, as previously described^24,26^, we assessed bulk activity in the parabrachial nucleus (PBN) as an indirect readout of ascending nociceptive information. We utilized *in vivo* fiber photometry to monitor calcium activity of glutamatergic neurons in the PBN during mechanical and cold stimulation before and after induction of MIA as well as after administration of either CBD3063 (10 mg/kg, i.p) or gabapentin (30 mg/kg, i.p) (**Fig. 3A**). As previously published, glutamatergic PBN neurons exhibited increased calcium responses to mechanical and thermal stimuli after MIA (**Fig. 2F-I**) compared to baseline (**Fig. 3B-E**), mirroring the increase in MIA-induced pain-like behaviors (**Fig. 2**). Both CBD3063 (**Fig. 3J-M**) and Gabapentin (**Fig. 3N-O**) reduced the MIA-evoked increases in calcium responses in response to mechanical and thermal stimulation (**Fig. 3R-U**). These results parallel our *in vivo* behavioral testing results (**Fig. 2**) and further validate the potential of CBD3063 as an OA pain relieving agent.

**Fig 3.**
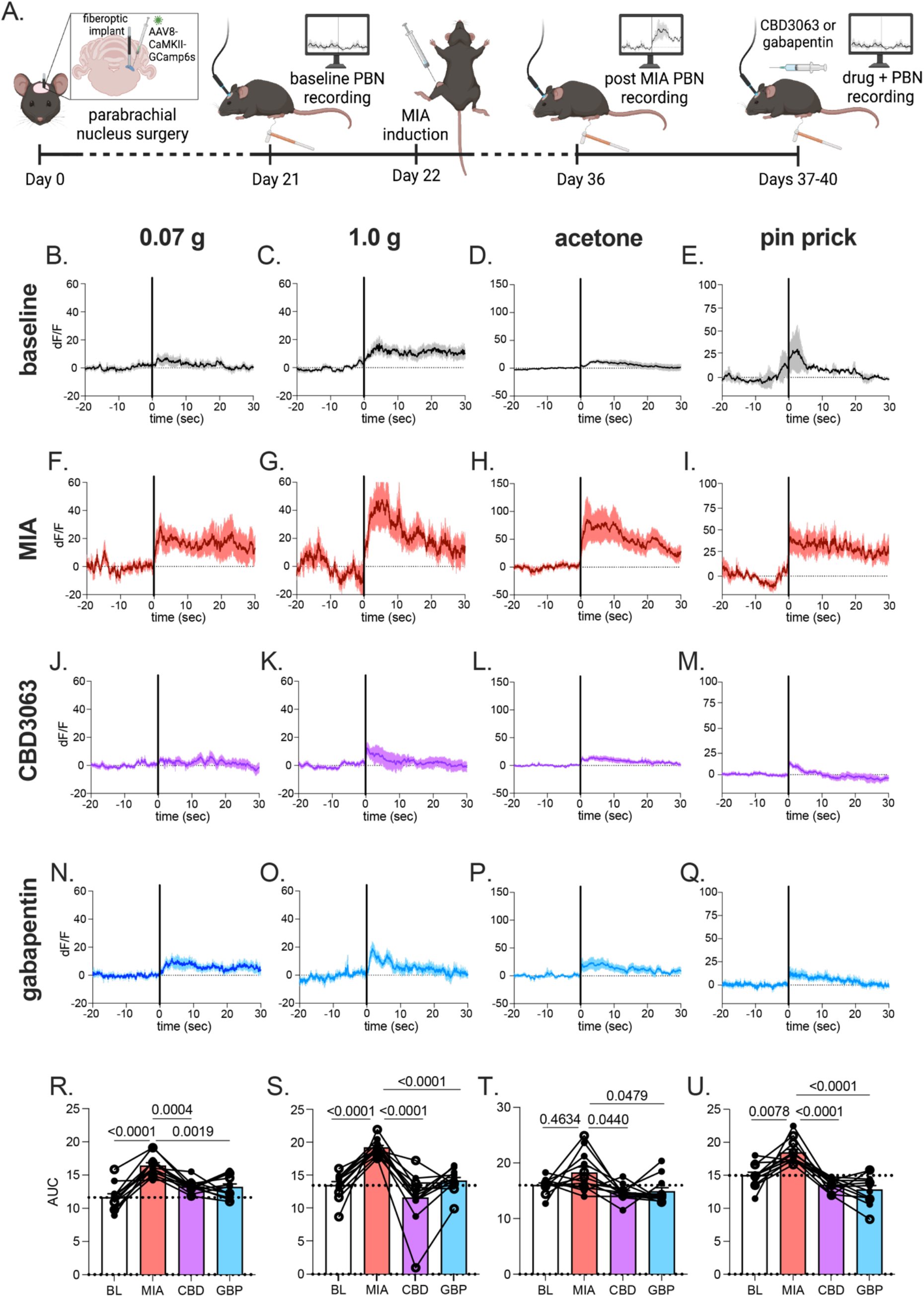
MIA increased glutamatergic PBN activity that was reduced by CDB3063 and gabapentin. (A) Schematic timeline for fiber photometry recording from glutamatergic PBN neurons. Averaged calcium transients in response to 0.07 g (B), 1.0 g von Frey filament (C), acetone droplet (D), and pin prick (E) at baseline. Average calcium response to 0.07 g (F), 1.0 g (G), acetone (H), and pin prick (I) two weeks after MIA-induction. Average calcium response to 0.07 g (J), 1.0 g (K), acetone (L), pin prick (M) after intraperitoneal injection of CBD3063 (10 mg/kg). Average calcium responses to 0.07 g (N), 1.0 g (O), acetone (P), and pin prick (Q) after intraperitoneal administration of gabapentin (30 mg/kg). Area under the curve summary data in response to 0.07 g filament (R), 0.1 g filament (S), acetone (T), and pin prick (U). Solid black vertical line at time 0 indicates application of stimulus (B-Q). Solid trace represents average response, shaded region indicates SEM. Results compared using one-way repeated measures ANOVA with Tukey posttest (R-U). N=12 animals, 8 females (filled circles)/4 males (open circles).

### CBD3063 decreases MIA-induced conditioned place aversion to mechanical stimuli

In addition to evaluating the analgesic efficacy of CBD3063 on the sensory discriminative aspects of pain, we also evaluated its ability to reduce the affective/motivational ongoing facets of pain. For this, we used the 2-chamber Conditioned Place Aversion (CPA) test (**Fig. 4A**), as previously reported^26^. Three weeks after the MIA-injury, rats were injected with either vehicle, CDB3063 (10 mg/kg) or gabapentin (30 mg/kg) and 1 hour later, the CPA-test was initiated (**Fig. 4A**). The dose and pre-treatment time were selected based on robust analgesic effects observed in the dose-response experiments for both compounds between 1-2 hours after injection, thereby covering the entire duration of the CPA-test (**Fig. 2**). At three weeks after MIA-injury, male (closed symbols) and female (open symbols) rats treated with vehicle spent significantly less time in the vF-conditioned chamber during the test (**Fig. 4B**), clearly demonstrating the aversion to the normally non-aversive stimuli induced by the MIA-injury. However, pre-administration of gabapentin (**Fig. 4C**) or CBD3063 (**Fig. 4D**) abolished the aversiveness of the mechanical stimuli. Quantification of the CPA-score revealed that both compounds significantly reduced the MIA-induced aversion compared to vehicle treatment (**Fig. 4E**).

**Fig 4.**
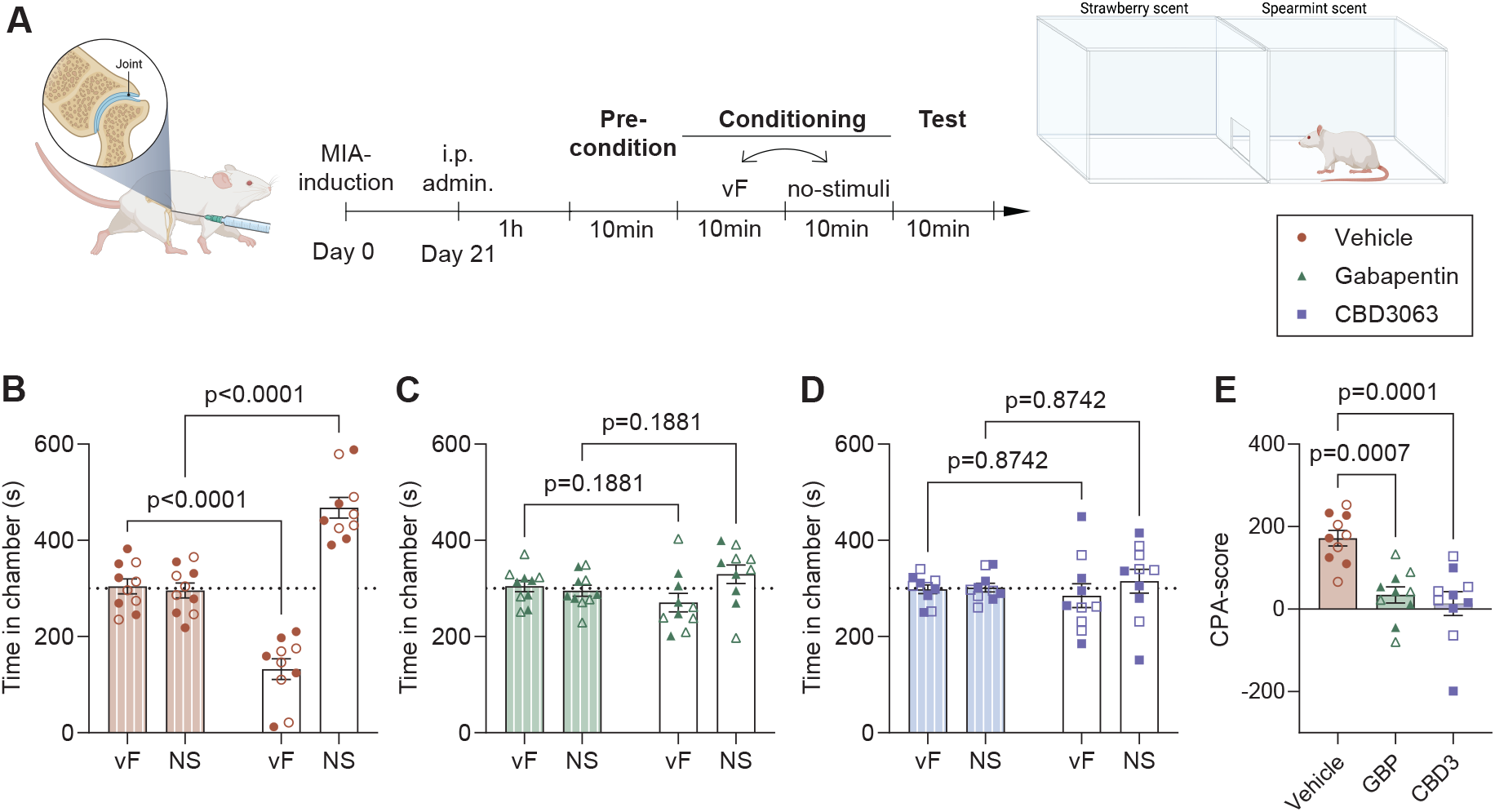
MIA-mediated behavioral aversion to mechanical stimulation is reduced by CBD3063 and gabapentin. (A) Schematic timeline of the two-chamber conditioned place aversion (CPA) test performed male and female rats 3 weeks after MIA-injury. 1h after injection of vehicle, gabapentin (30mg/kg) or CBD3063 (10mg/kg), the rat was exposed to 4*10 min consecutive sessions of; preconditioning, conditioning to each chamber, and test. Conditioning included one chamber conditioned to stimulation with a 10g vF-filament every 30second in one chamber, and no stimulation (NS) in the other. (B) Vehicle treated MIA-injured animals showed increased aversion to the VF-conditioned chamber during the test. (C)Injection of Gabapentin (30mg/kg) prevented the MIA-induced stimulus-aversion. (D) Injection of CBD3063 (10mg/kg) prevented the MIA-induced stimulus-aversion. (E) CBD3063 and gabapentin both decreased the MIA-induced aversion as seen by decreased CPA-scores when compared with vehicle. CPA score = time in VF-chamber during preconditioning – time in VF-chamber during test. For panel B-D; Results were compared using 2-way ANOVA and post-comparison with the Sidak’s multiple comparison test For panel E, results were compared using One-way ANOVA and Tukey’s post comparison test. N=10 across sex (5/5 pr sex). Females are displayed as open symbols, males as closed. Results are displayed as Mean ± S.E.M.

### Intraperitoneal administration of CBD3063 decreases MIA-induced weight bearing asymmetry

Weight bearing asymmetry is traditionally used as a surrogate marker for the pain experienced when weight is borne on the injured joint, and therefore a key non-evoked measure of how the joint injury affects and modifies the natural physical behavior^5^. During naïve circumstances, animals will carry equal proportions of weight on each hindleg (50%), but following induction of the MIA-injury, the animal will shift the majority of the weightbearing to the uninjured leg to compensate for the pain associated with weightbearing on the injured (ipsilateral) limb^2^.

We therefore wanted to clarify if CBD3063, in addition to improving the stimulus-induced hypersensitivity, also improved the pain-related weightbearing asymmetry. Approximately three weeks after injury (**Fig. 5A**), we confirmed that MIA-injury had caused a decrease (<50%) in weight borne on the injured leg (**Fig. 5B-C**, Post-MIA timepoint). Notably, we found that CBD3063 (10 mg/kg) and gabapentin (30 mg/kg) significantly improved the weight bearing asymmetry 2 hours after systemic administration in both male (**Fig. 5B)** and female rats (**Fig. 5C**).

**Fig 5.**
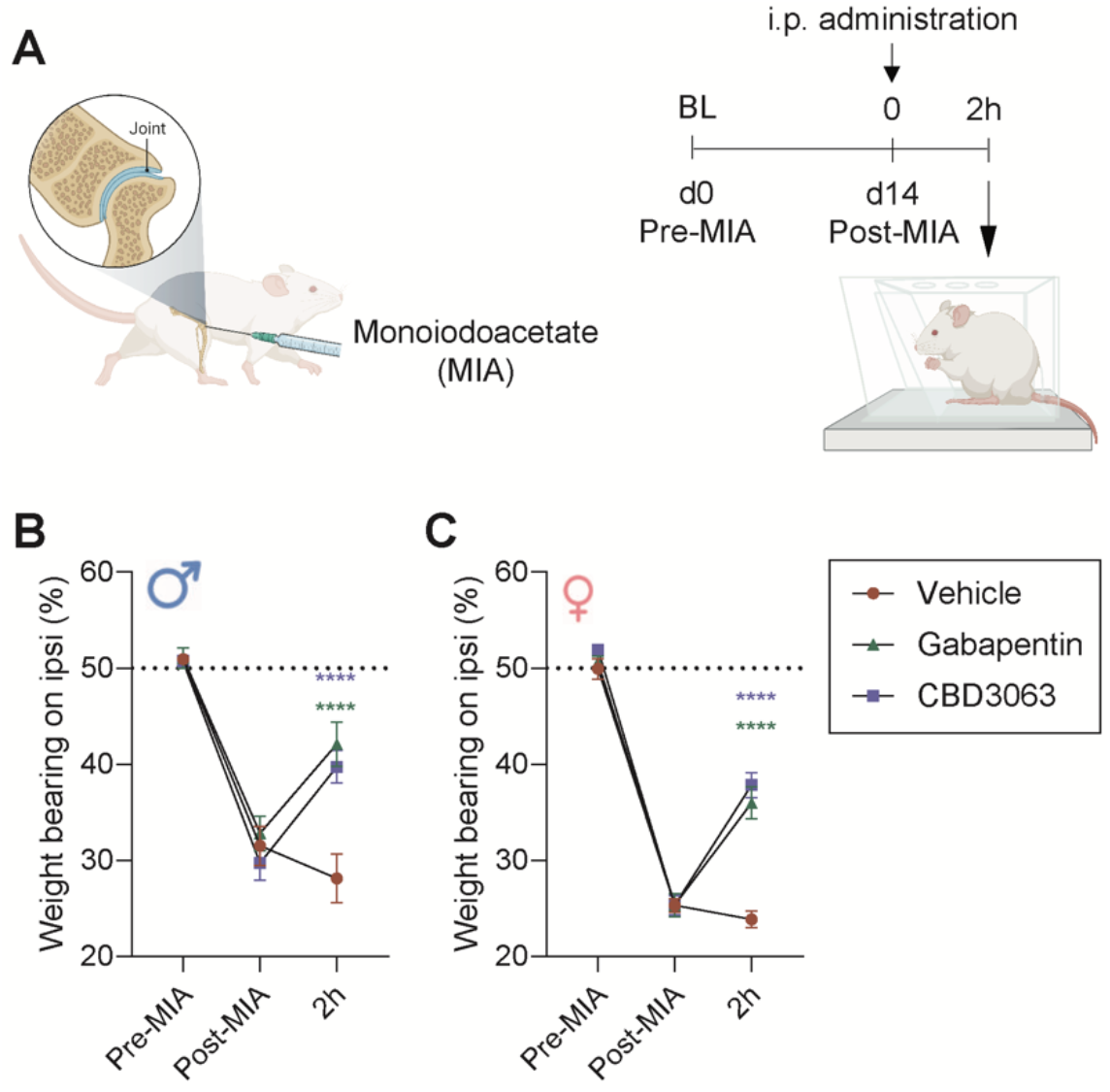
MIA-induced functional weight bearing deficits are improved by CBD3063. (A) Weight bearing distribution was assessed at baseline before MIA (pre-MIA), 3 weeks after MIA (post-MIA), and 2hours (2h) after injection of CBD3063 (10 mg/kg), gabapentin (30 mg/kg) or vehicle in male and female rats. MIA induced clear weight bearing asymmetry, seen as less weight born on the injured leg at the post-MIA, for both males (B) and females (C). The dotted line at 50% suggests the level for equal weight distribution between left and right hindleg. Treatment with CBD3063 and gabapentin produced significant increase in the proportion of weight born on the injured leg 2h after injection for both male and female rats. Results were compared using 2-way ANOVA and post-comparison with Tukey’s multiple comparison test, as suggested by; *P<0.05, **P<0.01, ***P<0.001. Error bars suggest Mean ± S.E.M. N=5 pr gender and treatment group.

## Discussion

Here, we present data demonstrating the potential of our first-in-class peptidomimetic, CBD3063, which indirectly regulates the trafficking of CaV2.2 calcium channels, as a novel and efficacious therapeutic agent for preclinical OA pain.

The N-type voltage-gated CaV2.2 calcium channel is a crucial target in the search for new pain treatments due to its significant role in pain signal transmission in the nervous system^12,44,52^. These channels, predominantly found in the nerve terminals of nociceptors synapsing in the spinal cord dorsal horn, are upregulated in chronic pain states^14,36,53^ and functionally contribute to nociception^22,49^. Further, genetic and pharmacological studies reveal the necessity of CaV2.2 for pain transmission^28,41^. Clinically, pharmacological inhibition of CaV2.2 has seen success with drugs like ziconotide, a synthetic peptide from the cone snail *Conus magus* venom, which is a potent and selective CaV2.2 blocker approved for severe chronic pain and administered intrathecally^42^. Further, gabapentinoids, such as gabapentin and pregabalin, indirectly reduce CaV2.2 activity by binding to the alpha-2-delta subunit of voltage-gated calcium channels, modulating their function^38^.

Despite these advancements, challenges persist, including significant side effects like motor dysfunction and dizziness due to the widespread distribution of these channels in the nervous system^20,30^. Consequently, targeted delivery methods like intrathecal administration are necessary but invasive^42^. In contrast, we believe our compound represents a superior strategy. By indirectly targeting CaV2.2 and regulating its trafficking, we aim to avoid many of the negative effects associated with the direct blockade of CaV2.2. We have repeatedly demonstrated that disrupting the interaction between CaV2.2 and CRMP2 with a short 15-amino acid peptide from CRMP2 (calcium channel binding domain 3, CBD3) is efficacious in reversing pain^7,23^. Further, we have continuously identified that disrupting CaV2.2–CRMP2 binding with tat-CBD3, myr-tat-CBD3 peptides, or CBD3063 does not affect memory, locomotion, nor anxiety/depression, and does not produce addictive behaviors^7,23,24^. Hogan and colleagues demonstrated sustained relief of nerve injury–induced neuropathic pain by an adenovirus expressing recombinant fluorescent CBD3 peptide (AAV-eGFP-CBD3), without evoking an anti-inflammatory response^54^.

Some limitations of our study should be noted. In addition to uncoupling the CaV2.2–CRMP2 interaction, the parent peptide, CBD3, also disrupts CRMP2 interactions with the N-methyl-D-aspartate receptor (NMDAR) NR2B subunit^6,8^ and enhances interactions with sodium calcium exchange 3 (NCX3)^10^ to confer neuroprotection in preclinical models. Whether CBD3063 also engages these mechanisms to bring about the anti-hyperalgesia is not known. This study does not address the cellular site of action of CBD3063’s anti-hyperalgesic efficacy. However, since CRMP2 and CaV2.2 are expressed in multiple sites along the pain neuraxis, CBD3063 disruption of their interaction is likely to be beneficial at all sites. The study did not address if CBD3063 treatment could be disease modifying.

Markedly, in contrast to neuropathic pain, the use of CaV2.2 directed agents for the treatment of arthritis-related pain has been limited both preclinically and clinically. Our data shows, for the first time, that MIA increases CaV2.2 protein in the spinal cord irrespective of sex. While ample data implicates CaV2.2 as a drug target for neuropathic pain, less data substantiates this key channel for the treatment of OA pain. Gabapentinoids have demonstrated efficacy in some rodent models of OA^29^ and human OA clinical studies^19,45^.

Further, gabapentin improved weight bearing^31^ and owner-perceived mobility impairment^25^ in dogs and cats with OA, respectively. Here, we demonstrate that targeting CaV2.2 reduces both sensory-discriminative and affective-motivational aspects of pain in a rodent model of OA. Further, our use of *in vivo* fiber photometry reveals CBD3063 effectively diminished the transmission of nociceptive information from the spinal cord to the brain via the spino-parabrachio-amygdaloid pathway. This is in line with findings which reported higher levels of the neuropeptides – substance P (SP) and calcitonin gene-related peptide (CGRP) – in the spinal cord in a rat model of OA^21^. Together, these results suggest that future studies should continue to evaluate CaV2.2-directed pharmacotherapeutics for the treatment of OA pain.

## Acknowledgements

This work was supported was supported by National Institutes of Health (NIH) awards F32NS128392 (to HNA), K00NS124190 (to TSN), and RF1NS131165, R61NS126026, R01NS120663 (to RK). Additionally, this work was supported by a Development Grant from the American Neuromuscular Foundation (to TSN).

## Notes

**Conflict-of-interest:** R. Khanna is the co-founder of Regulonix LLC, a company developing non-opioids drugs for chronic pain. The rest of the authors have declared no conflict of interest.

### Competing Interest Statement

R. Khanna is the co-founder of Regulonix LLC, a company developing non-opioids drugs for chronic pain.

