## Supplementary Table 1 for "Uncoupling the CRMP2-Ca_V_2.2 interaction reduces pain-like behavior in a preclinical osteoarthritis model"

**Supplementary table, S1. Statistical analysis**

| **Figure** | **Analysis** | **Outcome** | **Post-test** |
| --- | --- | --- | --- |
| **Fig 1. MIA enhances the presynaptic fraction of Ca_V_2.2.** | | | |
| Fig 1C – Ca_V_2.2 | Unpaired two-tailed t test, Vehicle vs. MIA | P=0.0035 | N/A |
| Fig 1C – CRMP2 | Unpaired two-tailed t test, Vehicle vs. MIA | P=0.5858 | N/A |
| **Fig 2. CBD3063 decreases mechanical and cold allodynia in rats with MIA-induced pain-like behavior** | | | |
| Fig 2B. Mechanical allodynia - males | Two-way RM ANOVA, time*treatment | F_time_(8,72)=24.14, P<0.0001  F_treatment_(2,9)=70.32, P<0.0001  F_time*treatment_(16,72)=2.948, P=0.0009 | Tukey  N=4 |
| Fig 2C. Mechanical allodynia - females | Two-way RM ANOVA, time*treatment | F_time_(8,72)=29.35, P<0.0001  F_treatment_(2,9)=6.479, P=0.0181  F_time*treatment_(16,72)=2.949, P=0.0009 | Tukey  N=4 |
| Fig 2D. Mechanical allodynia - AUC | Two-way RM ANOVA, sex*treatment | F_sex_ (1,18)=0.136, P=0.72 (NS)  F_treatment_(2,18)=34.07, P<0.0001  F_sex*treatment_(2,18)=0.42, P=0.66 (NS) | Tukey  N=4 |
| Fig 2E. Cold allodynia - males | Two-way RM ANOVA, time*treatment | F_time_(4,36)=101.7, P<0.0001  F_treatment_(2,9)=21.56, P=0.0004  F_time*treatment_(8,36)=13.78, P<0.0001 | Tukey  N=4 |
| Fig 2F. Cold allodynia - females | Two-way RM ANOVA, time*treatment | F_time_(4,36)=75.25, P<0.0001  F_treatment_(2,9)=24.10, P=0.0002  F_time*treatment_(8,36)=9.157, P<0.0001 | Tukey  N=4 |
| Fig 2G. Cold allodynia - AUC | Two-way RM ANOVA, sex*treatment | F_sex_ (1,18)=18.14, P=0.005  F_treatment_(2,18)=61.53, P<0.0001  F_sex*treatment_(2,18)=2.61, P=0.10 (NS) | Tukey  N=4 |
| **Fig. 3 CDB3063 reduces glutamatergic PBN activity** | | | |
| Fig 3R. 0.07 g filament - AUC | One-way RM ANOVA | F(2.612, 28.73)=22.44, P<0.0001 | Tukey  N=12 |
| Fig 3S. 1.0 g filament - AUC | One-way RM ANOVA | F(1.979, 21.77)=27.47, P<0.0001 | Tukey, N=12 |
| Fig 3T. acetone – AUC | One-way RM ANOVA | F(2.162,23.79)=5.512, P=0.0094 | Tukey, N=12 |
| Fig 3U. pin prick – AUC | One-way RM ANOVA | F(2.429,26.72)=27.46, P<0.0001 | Tukey, N=12 |
| **Fig 4. MIA-mediated behavioral aversion to mechanical stimulation is reduced by CBD3063.** | | | |
| Fig 4B – vehicle treatment, time in chamber | Two-way RM ANOVA, test-phase*chamber | F_chamber_ (1,18) = 50.87, P<0.0001  F_test phase*chamber_ (1,18) = 165.4, P<0.0001 | Sidak  N=10  across sex |
| Fig 4C – Gabapentin treatment, time in chamber | Two-way RM ANOVA, test-phase*chamber | F_chamber_ (1,18) = 1.9, P=0.18 (NS)  F_test phase*chamber_ (1,18) = 6.055, P=0.0242 | Sidak  N=10 across sex |
| Fig 4D – CBD3063 treatment, time in chamber | Two-way RM ANOVA, test-phase*chamber | F_chamber_ (1,18) = 1.02, P=0.33 (NS)  F_test phase*chamber_ (1,18) = 0.44, P=0.51 (NS) | Sidak  N=10 across sex |
| Fig 4E – CPA score | One-way ANOVA | F (2,27)=14.09, P<0.0001 | Tukey  N=10 across sex |
| **Fig 5. Intra peritoneal administration of CBD3063 decreases MIA-induced weight bearing asymmetry.** | | | |
| Fig 5B –weight bearing males | Two-way RM ANOVA, time*treatment | F_time_(2,24)=92.34, P<0.0001  F_treatment_(2,12)=8.616, P=0.0048  F_time*treatment_(4,24)=5.775, P=0.0021 | Tukey  N=5 |
| Fig 5C – weight bearing females | Two-way RM ANOVA, time*treatment | F_time_(2,24)=682.5, P<0.0001  F_treatment_(2,12)=13.82, P=0.0008  F_time*treatment_(4,24)=22.83, P<0.0001 | Tukey  N=5 |
